# A predictive model of rats’ calorie intake as a function of diet energy density

**DOI:** 10.1101/184085

**Authors:** Rahmatollah Beheshti, Yada Treesukosol, Takeru Igusa, Timothy H. Moran

**Author notes:** Corresponding author. Address: Room W3510, Bloomberg School of Public Health, 615 N. Wolfe Street, Baltimore, MD 21205. **Author contributions:** RB designed and conducted research, analyzed the data, and wrote the manuscript. TM and TI provided supervision for the model development and implementation and provided critical input to the manuscript draft. YT collected original datasets in separate studies and provided critical input to the design of the research, drafted portions of the manuscript and provide critical input to the manuscript draft. All authors read and approved the final manuscript.

## Abstract

Easy access to high-energy food has been linked to high rates of obesity in the world. Understanding the way that access to palatable (high fat or high calorie) food can lead to overconsumption is essential for both preventing and treating obesity. Although the body of studies focused on the effects of high energy diets is growing, our understanding of how different factors contribute to food choices is not complete. In this study, we present a mathematical model that is able to predict rats’ calorie intake to a high-energy diet based on their ingestive behavior to a standard chow diet. Specifically, we propose an equation that describes the relation between the body weight (W), energy density (E), time elapsed from the start of diet (T), and daily calorie intake (C). We tested our model on two independent data sets. Our results show that the suggested model is able to predict the calorie intake patterns with high accuracy. Additionally, the only free parameter of our proposed equation (ρ), which is unique to each animal, has a strong correlation with their calorie intake and weight gain. Additionally, we discuss the relevance of our derived parameter in the context of measuring reward sensitivity in reinforcement learning based studies.

## Introduction

Through the course of history, the human body has evolved to survive times of food deprivation. Additionally, through innate and learned responses, we generally find energy-dense foods the most palatable, and as such have learned which cues indicate the presence of those foods (33). Changes in agriculture and the food industry during the past century have provided unprecedented access to cheap, energy-dense foods. It has been argued by many scholars that one of the major contributing factors to the worldwide obesity epidemic has been easy access to highly palatable, energy-dense foods (24). In such a situation, understanding the role of rewarding aspects of food intake, including its appetitive and consummatory aspects, can help us find more efficient ways to prevent and mitigate the obesity epidemic. Despite the robust correlation observed between the functionality of the food reward system (i.e. the neural pathways that play a role in food wanting, liking and reinforcement) and obesity, our knowledge about the rewarding aspect of food is incomplete, mainly due to its complex nature. Mathematical and computational approaches to model different contributing factors of food decision making is one way to understand the complexities of these components, which has recently received much attention. While it is tough to control and study all of these contributing factors to food behaviors in human settings, animal models can minimize some of the complexities of human studies and thus provide valuable insights into understanding of these interactions. Developing models that can describe food intake patterns in animals (without including the connection to the rewarding aspects) has been the subject of many works. For instance, Lusseau et al. presented a mathematical model for describing the food intake patterns of mice that undergo calorie restrictions using a Markov model by setting the probabilities of transition between the states of the Markov model (representing shifting across different physiological states) (21). In addition to modeling food intake patterns, other studies have presented mathematical models for studying energy metabolism in animals (3, 9, 18). For instance, Guo et al. proposed several models to predict changes in body weights and energy expenditure of mice in different conditions (10, 11). Also, from a different point of view, Jacquier et. al. presented a series of differential equations to model the hormonal regulation of food intake in rats (13–15).

While various types of models have been proposed for describing different aspects of food intake in animals, a model that links food intake patterns and energy density of food does not exist. High energy foods—like those high in sugar or fat—are generally considered more palatable and more rewarding. In this paper, we present a mathematical model that is able to predict the amount of calorie intake in rats as a function of the energy density of the diet. Our proposed model consists of an equation that includes four main variables: body weight (*W*), energy density of the diet (*E*), elapsed time since the beginning of the current diet (*T*) and finally a parameter *ρ* (rho) peculiar to each rat.

### Materials and methods

We have used several existing models of food intake as the basis of developing our model. For inferring the level of perceived reward (R) in rodents, one objective measure that is commonly used in other works is licking rate when a liquid meal is consumed. It has been shown that licking rate for each individual animal follows a Weibull function in the form of *y=Aexp[-(Bt)^c^*], where *A, B,* and *C* are the parameters unique to each animal, and *t* is the time passed after the food was presented (6, 25). Similar exponential equations have been shown to be capable of modeling the relationship between the cumulative amount of calorie intake (C) and the meal time (23). Based on these two sets of formulations, the relation between the cumulative calorie intake (C) and reward (R) will follow a form of decreasing exponential curve with a form that is schematically shown in Fig. 1. As eating commences (or more precisely shortly after the beginning), the amount of reward is at the highest point. As more food is consumed, the reward also gradually decreases until it reaches zero.

**Fig. 1.**
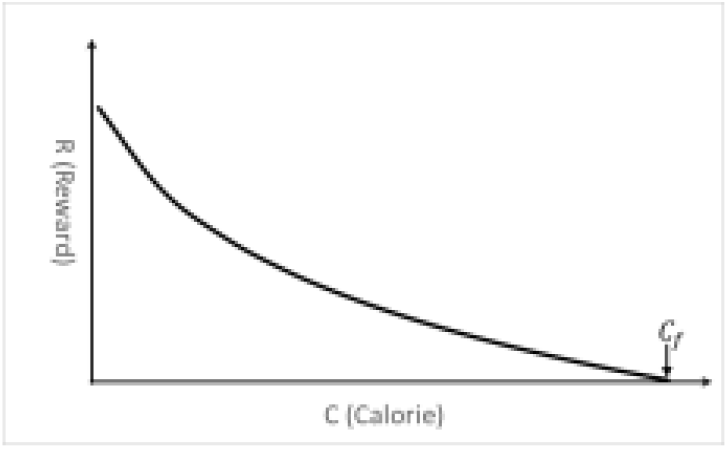
A simple representation of the relation between the amount of the perceived reward and calorie intake (C) in a meal. The reward curve follows a logarithmic function, which becomes zero at Calorie=C_f_.

For *C>0,* this relation can be simplified with a logarithmic function. More details about this approximation are provided in Appendix. Such a logarithmic relation between *C* and *R* can be shown in a function that has the form of:

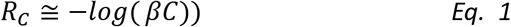

Where, *R_C_* shows the amount of perceived reward after consuming *C* calories, and *β>0* is a constant multiplier. Here, reward refers to the palatability of food as inferred by the licking rate. To be able to describe different calorie intake patterns for different individual animals using this equation, the multiplier has to be specific to each animal. To determine the factors that affect the value of *β*, we consider a set of major elements that are known to influence the rewarding aspects of food intake and therefore β. Many studies have reported a positive effect of body weight (W) on the amount of experienced reward (7, 8). We also know that the amount of perceived reward has a direct relation to the energy density of the diet (E) (4, 29), and an inverse relation with the time elapsed from the first time that a new diet was introduced (T) (28). As *-log(x)* is a decreasing function, to include these variables with similar effects we can replace *β with a new formula, such as:*

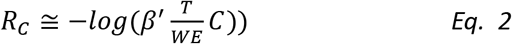

Where, β’>0 is a new constant multiplier specific to each individual animal. In this suggested formulation, higher values of *T* would make the slope of the curve in Fig. 1 steeper, while higher values of *W* or *E* would have an opposite effect. Based on a similar logic, Eq. 2 can be considered in other ways as well. More generally, W, E, and *T*can be replaced by *f(W), f(E)* and *f(T),* while *f* is a strictly increasing function, such that a similar relation between these variables is held. In the next section, we discuss the specific functions that worked best for this equation. Based on our model, eating ceases when the value of perceived reward is equal to zero. Both homeostatic and hedonic factors affect reaching this stopping point, but collectively the combination of all of the different factors makes calorie intake rewarding at first, and when the calorie intake is not appealing anymore the reward value equals zero. This stopping point is shown in Fig. 1 by *C_f_* (subscript *f* for final). As *C = C_f_* is the point where *R*_*c*_ reaches to zero, from Eq. 2 we will have:

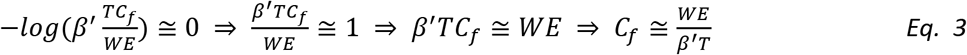

In other words, we can expect that the amount of calories that an animal consumes can be predicted based on the values of *W, E, T* and *β’*. For simplicity, we can bring *β’* to the numerator, and instead use a new parameter *ρ:*

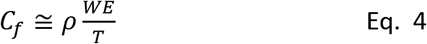

The process described so far was used to find a starting point for our search for finding a logical mathematical model that can describe the relation between calorie intake and energy density of food. To see whether a formula in this form can be applied to the actual cases, and also determine which *f* functions *(f(W), f(E)* and *f(T))* are the best, we have used two data sets from two independent groups of rats. In the following, these two datasets are briefly introduced, before presenting the best form of Eq. 4 that was found based on these datasets.

## Data

<u>Study-1:</u> The first dataset was related to a group of (n=16) male Sprague Dawley (Harlan) rats ∼275-300 g at the age of 60-70 postnatal days. Following seven days of habituation to the laboratory environment and maintenance on a chow diet with an energy density of 3.1 kcal/g (2018 Teklad, Harlan), the animals were assigned to two groups of eight. One group was maintained on the chow diet, and the other group was switched to a high-energy (HE) diet with an energy-density of 4.73 kcal/g (D12451, Research Diets) for 42 consecutive days. At this point, the diet of the HE group was switched back to the original chow diet for an additional seven consecutive days of recovery. Additional details about this study are also provided in the original article (32).

<u>Study-2:</u> The second dataset was collected through another similar experiment using (n=20) rats of the same type, and similar diets. This time, after 14 days of habituation with the chow diet (3.1 kcal/g), half of the rats received HE diet (4.3 kcal/g) for 14 days.

We used the data from the rats that received HE diet for the purpose of training and testing of our model. In the case of Study-1, the data from both the initial chow diet and the recovery period were used for training our mathematical model; and in Study-2, only the data from the initial chow diet was used for training. The reason for considering the recovery period for the Study-1 was that the existing data for the initial chow diet period in the first experiment was insufficient (only four days for each rat), and this was not enough to properly train our model. Training refers to finding the best values of *ρ* by fitting (calibrating) the equation to actual data. Training was performed using nonlinear regression analysis *(fitnlm* and associated functions) in the MATLAB software. It returns a nonlinear model representing a least-squares fit of the response to the data. After training the model using the chow diet data, it was then tested on the data from the HE diet period. The model was trained and tested separately for each individual rat, using the available data.

## Results

Before training our model on the rat datasets (determining the best value of *ρ* for each rat), we needed first to find the best form of Eq. 4. We tested a set of common mathematical functions that can be used for the variables *W, E* and *T* in Eq. 4, with arbitrary (synthetic) values of *ρ*. Specifically, we tested a range of power functions (x^c^ (c is a constant value)) and exponential functions (e^x^ and log(x)), and found that an equation in the following form can generate the closest patterns to the available data:

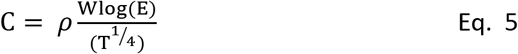

At this stage, we wanted to find a form of Eq. 4 that shows the highest potential for matching to the data. A similar procedure for finding the best equation to analyze food intake patterns has been used by others (16). More details about this procedure are provided in the Appendix. We then fitted Eq. 5 to the training data as described earlier, to evaluate the performance of this candidate equation. During this process, using only chow data, the best values of *ρ* were determined for each rat. The trained model was then tested by using the model to predict the amount of calorie intake during the period of HE diet. Having the values for ρ, *W, E,* and T, we used Eq. 5 to predict calorie intakes (C). The results of testing our model are shown in Fig. 2 (Study-1 rats) and Fig. 3 (Study-2 rats). Fig. 2 illustrates the comparison between the actual calorie intake of the eight rats of Study-1 and the amount of calorie intake predicted by our model. These amounts of calorie intake refer to the period of receiving HE diet (the testing period). Each chart shows the daily calorie intake of one rat for a period of 42 consecutive days *(T=1-42).* During this period, energy density (E) of the rats’ diet was equal to 4.3 kcal/g, and body weights of rats (*W)* were also available from the data.

**Fig. 2.**
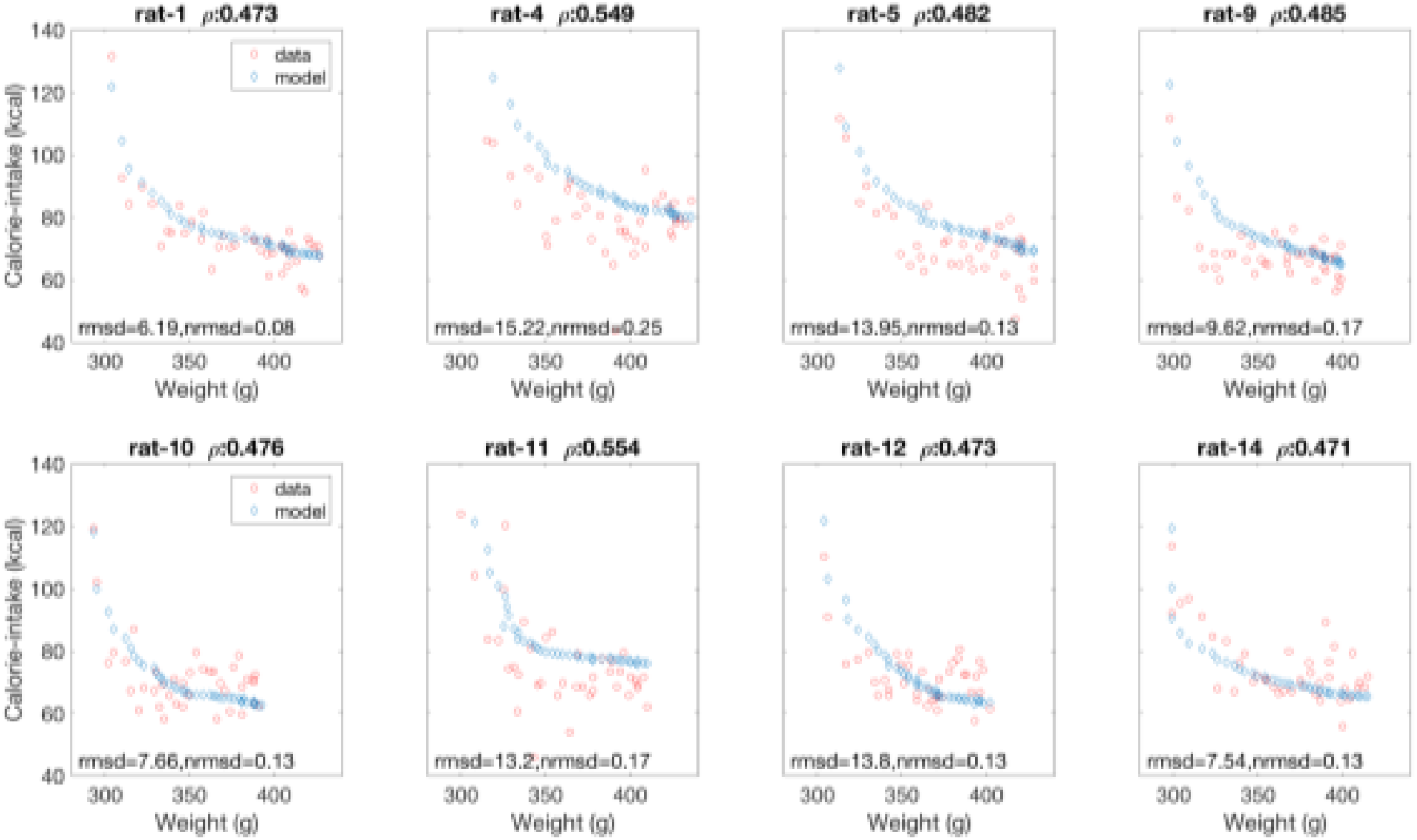
Comparison between the actual daily calorie intake of eight rats from the Study-1 (red circles) and the predicted calorie intake by our model (blue diamonds) during the 42 days of high-energy diet presentation. Rat ID and the value of ρ (rho) for each rat are shown on top. rmsd: root-mean-square deviation; nrmsd: normalized root-mean-square deviation.

Similarly, Fig. 3 shows the comparisons between the actual calorie intakes of the ten rats of Study-2 during the 14 days of receiving HE diet and our model’s predictions.

**Fig. 3.**
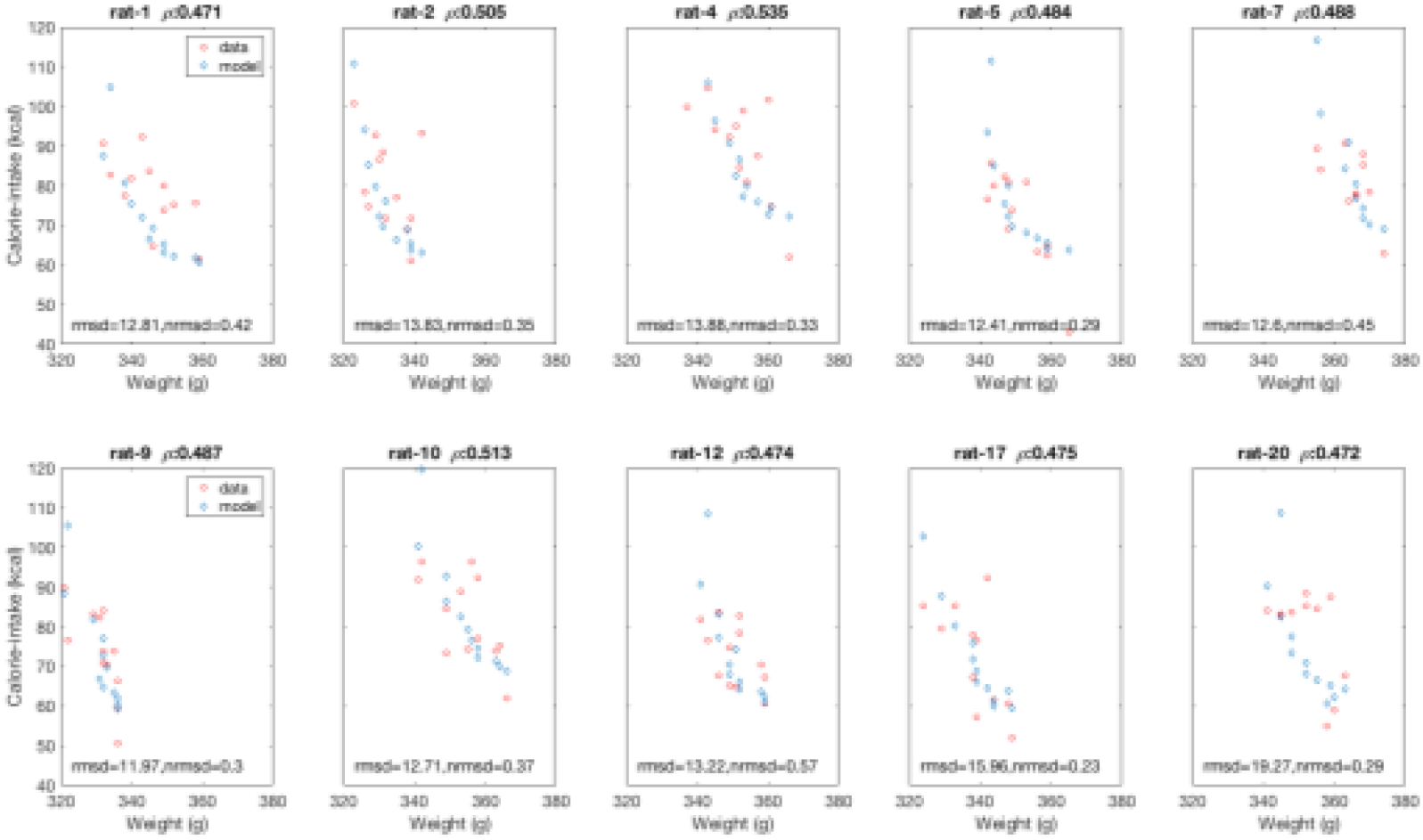
Comparison between the actual calorie intake of the ten rats of the Study-2 (red circles) and the predicted ones by our model (blue diamonds) during the 14 days of receiving high-energy diet. Rat ID and ρ (rho) value for each rat are shown on top. rmsd: root-mean-square deviation, nrmsd: normalized root-mean-square deviation.

To explore the extent that the value of *ρ*—which is specifically assigned to each rat using the training data—is capable of predicting the calorie intake and weight gain of each rat, we ran a set of linear regression analyses. As mentioned earlier, the *ρ* values were calculated (trained) using the chow diet data, and not the HE diet. Fig. 4A and Fig. 4B show the correlation between *ρ* and the average amount of daily calorie intake during the period of receiving HE diet.

**Fig. 4.**
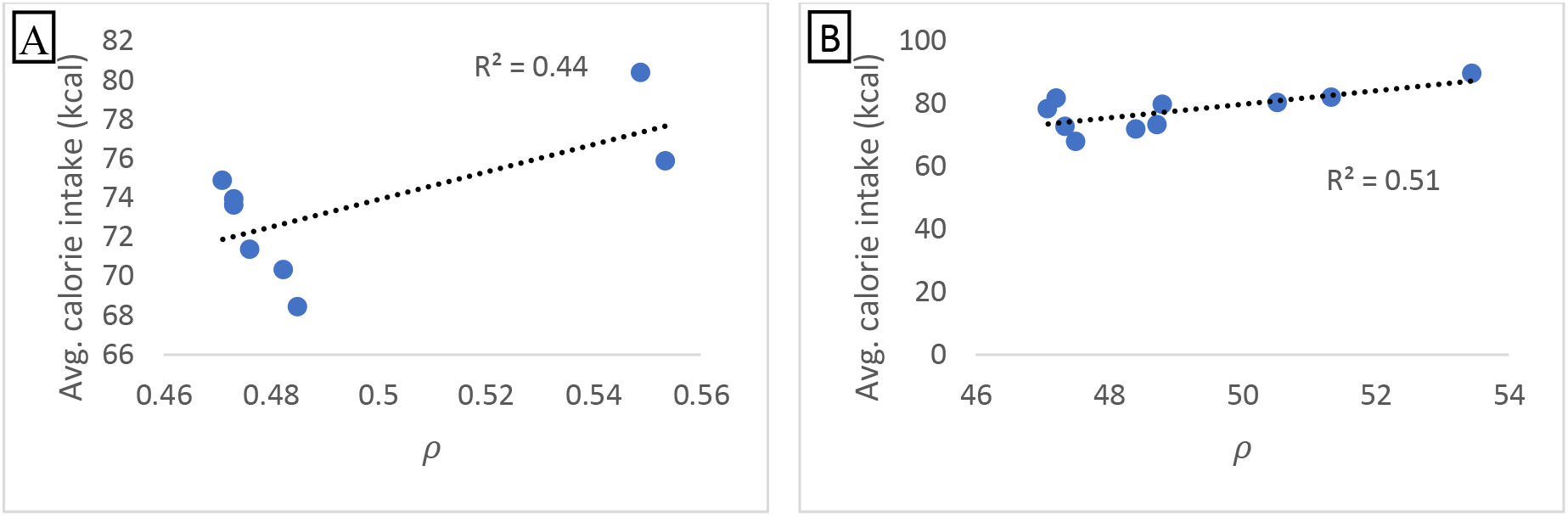
The correlation between the values of ρ (rho) for each rat and the average daily calorie intake during the period of high-energy diet. (A) Eight rats of Study-1, (B) 10 rats of Study-2.

Similarly, the correlation between the percentage of body weight gain and the values of *ρ* are shown in Fig. 5. The body weight of animals at the beginning of experiments was considered as 100%, and the reported percentages in Fig. 5 show the total increase by the end of the HE diet period. For instance, 44% indicates that the overall body weight of an animal reached to 144% by the end of the HE diet period.

**Fig. 5.**
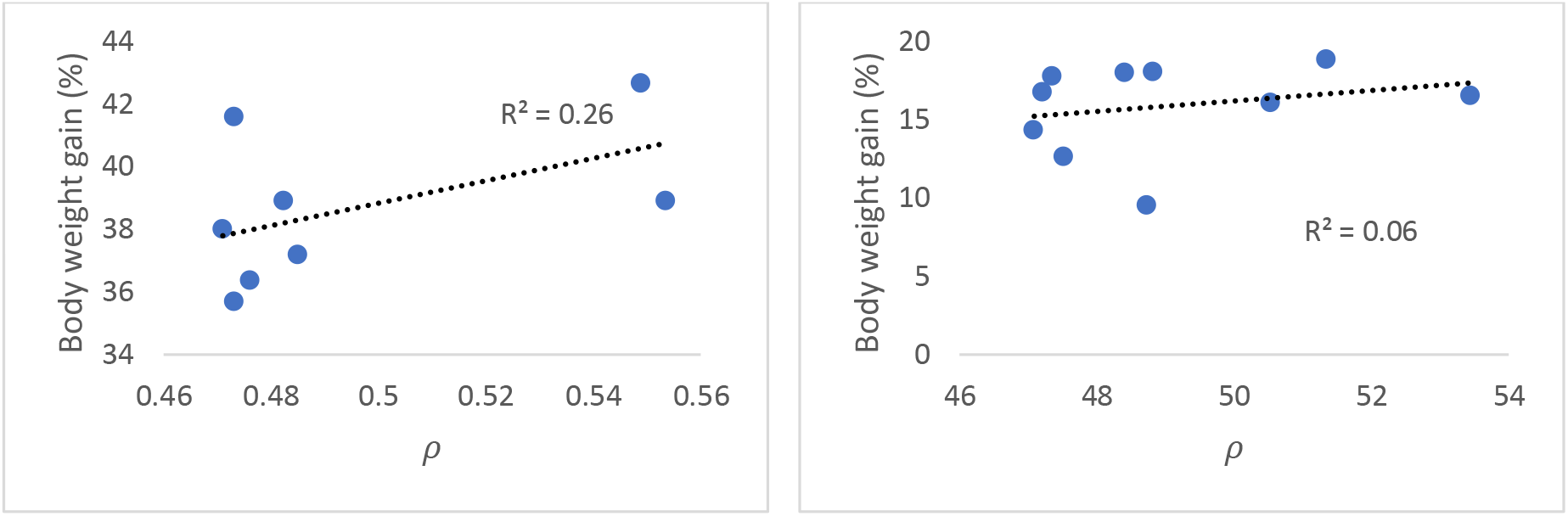
The correlation between the percentage of body weight gain by the end of the high-energy diet period and the values of ρ for each rat in (A) Study-1, (B) Study-2.

## Discussion

Our model presents a simple equation that can predict the amount of calorie intake of rats, based on their body weight, the energy density of the food and the length of the period that the food was presented. These variables have been reported in various studies as the major factors that affect the food intake of rodents. For instance, in a recent study, the set of the main covariates of food intake that were considered for modeling the effects of calorie restriction on the behavioral phenotype of a group of mice included several variations of the time elapsed from the beginning of each diet change and body weight of animals (21).

Our model meets the four criteria in the so-called “first principles modeling approach” that have been identified by others (1, 23, 31) for a candidate model to constitute a good model of food intake behaviors. Specifically, our model has a self-consistent theoretical basis that described earlier. Additionally, it is a simple while not too simple model (one of the criteria). In fact, a major strength of our proposed model is that it has only one free parameter that needs to be set. This aspect leads to a parsimonious model, which is generally considered as a desirable property of mathematical models (22).

The results obtained from testing our model on the two independent datasets showed that our model can predict the calorie intake patterns of rats with relatively high accuracy. To formally measure the distance between the actual datasets and predicted patterns by our model, root-mean-square deviation (rmsd), and normalized root-mean-square deviation (nrmsd) between these two datasets are shown in Fig. 2 and Fig.3 for each rat. Both rmsd and nrmsd are common ways of measuring the accuracy of different models. The range of values of rmsd was between 6 and 19, showing that our model’s predictions were close to the actual calorie intake patterns. Additionally, the values for normalized rmsd measurement were between 0.08 and 0. 57 (with an average of 0.15 for Study-1 and 0.36 for Study-2).

Regression analysis was used to test if the values of *ρ* could significantly predict the average amount of calorie intake during the HE diet. The results of the regression indicated that the predictor explained 44% and 51% of the variance in two studies (R^2^=0.44, p=0.05; R^2^=0.51, p=0.02). Similarly, the regression analysis on *ρ* values and the rate of weight gain by the end of HE diet showed that *ρ* values explained 26% and 59% of the variance respectively (R^2^=0.26, p=0.05; R^2^=0.59, p=0.02). These results show that the differences in the value of parameter *ρ* can predict different calorie intake levels across dietary shifts. As each rat’s value of *ρ* is unique, this is in line with other studies that reported the role of genetic background in establishing individual differences in the responses to the changes in diet composition, energy availability or both (20). An immediate extension of our work would be applying our model to two genetically different groups of rats (e.g. rats with diet induced obesity (DIO) and diet resistant (DR) rats), and measure the differences between the *ρ* values of the two groups. We expect that a significant difference should exist between such groups.

Food with higher energy densities, such as high sugar and high-fat foods, are generally considered more palatable and more rewarding to humans and non-human animals. As our model is specifically designed to predict the calorie intake of animals when a diet with a different energy density is offered, it directly relates to the studies of the rewarding aspects of food intake. Among these are the studies based on the principles of reinforcement learning. Specifically, the behavior and meaning of the parameter *ρ* in our model closely relates to the so-called “reward sensitivity” parameter in the reinforcement learning models of food choice, which has many applications in different studies of rewarding aspects of food intake (17). Reward sensitivity is a variation in the primary sensitivity to rewards. Reduction in reward sensitivity is possibly the closest behavioral equivalent to the notion of a reduction in consummatory pleasure. Multiple studies have linked body weight changes to “reward sensitivity” and “learning rate” of individuals (5, 8, 17, 30), which are the two standard measures used in developing models of reinforcement learning. These two parameters are commonly calculated using maximum likelihood estimation methods on the datasets collected in probabilistic reward tasks experiments (12). Because of the way that *ρ* is defined in our model, it encodes a similar meaning as the reward sensitivity parameter. As collecting data using traditional ways—i.e. from iterative probabilistic reward tasks— is challenging, our model can potentially serve as a simpler way to calculate the reward sensitivity parameter. Investigating this hypothesis (ρ indicates the reward sensitivity) would be the subject of another study.

In addition to the palatability of food (preference value for a particular food), it has been reported that food variety (availability of different types of food) also has an incremental and distinguishable effect on food consumption and meal parameters (26, 27). In our model, the time elapsed after the change in the diet (T in our model) closely relates to the food variety factor. Similar to the effects of presenting the same food vs. a new one (as captured by T), it has been shown that food intake increases as the variety of presented food increases. This also relates to the *sensory-specific satiety,* as extensively studied in the literature (2).

The generated patterns of food intake by our model match many of the patterns reported in other studies of food intakes in rodents. One pattern that we observed in our simulated results was that after switching to a high energy diet, the generated calorie intake patterns initially increased significantly, and then decreased across subsequent days. In addition to matching to our actual data, this is also consistent with the other studies, which suggest that increased positive signals from oral stimulation and dampened inhibitory feedback are most robust across the first few days upon high-energy diet presentation (32). Moreover, we ran another experiment to use our model to predict body weight changes of Study-2 rats during the recovery period (using W = C.T^¼^. (log(E). ρ)^-1^ derived from Eq. 5). This experiment is presented in Appendix. We observed that as the calorie intake of rats (model’s input) decreased, simulated body weights by our model also plateaued. These patterns match the same patterns reported by Levin et al. (19, 20). They observed that rats that gained weight during high energy diet presentation return to a so-called “set point” after representation of a chow diet.

Lastly, our model can also be tested on other types of rodents or animals. While testing our model on one data set (using independent training and test data) could suffice the goal of validating our model, testing our model on a second dataset let us have increased confidence in the model. Also, our model only considers the total daily food intake of rats at this point, and implicitly assumes that all of the daily food intake is consumed in only one meal. The model can be extended in the future to also simulate meal patterns, such as the meal size and frequency of meals.

